# Insights into Cellular Evolution: Temporal Deep Learning Models and Analysis for Cell Image Classification

**DOI:** 10.1101/2024.03.11.584308

**Authors:** Xinran Zhao, Alexander Ruys de Perez, Elena S. Dimitrova, Melissa Kemp, Paul E. Anderson

## Abstract

Understanding the temporal evolution of cells poses a significant challenge in developmental biology. This study embarks on a comparative analysis of various machine-learning techniques to classify sequences of cell colony images, thereby aiming to capture dynamic transitions of cellular states. Utilizing transfer learning with advanced classification networks, we achieved high accuracy in single-timestamp image categorization. We introduce temporal models—LSTM, R-Transformer, and ViViT—to explore the effectiveness of integrating temporal features in classification, comparing their performance against non-temporal models. This research benchmarks various machine learning approaches in understanding cellular dynamics, setting a foundation for future studies to enhance our understanding of cellular developments with computational methods, contributing significantly to biological research advancements.

## II. Introduction

Recent advancements in developmental biology and bioengineering have paved the way for groundbreaking approaches in regenerative medicine and cellular therapies, particularly through the applications of stem cells [1]. The study of morphogenesis reveals the complex processes by which cell populations coordinate to form intricate structures from a single cell [2]. Human induced pluripotent stem cells (hiPSCs) have emerged as an essential resource for investigating the processes of cellular organization, differentiation, and assembly into tissues [3].

Building on recent advancements in developmental biology, bioengineering, and the utility of hiPSCs, our research is fundamentally aimed at exploring the capacity of computational approaches to accurately capture the temporal dynamics of stem cell morphogenesis. We concentrate on cellular structures as represented in microscope images captured over a 24-hour period of hiPSC evolution. At the heart of our investigation is the challenge of classifying cell colony images across these timestamps, a task that not only tests the limits of current computational biology methods but also seeks to illuminate the complex process of cellular development and differentiation.

By analyzing models trained on single-timestamp images, we explore network characteristics over time to identify temporal patterns, validating the presence of temporal context within our dataset. This leads us to assess the effectiveness of advanced sequential models which incorporate these temporal features. Then by conducting a thorough comparative analysis of temporal models like Long Short Term Memory Networks (LSTM), R-Transformers, and the Video Vision Transformer (ViViT), we assess these temporal models’ performance compared with non-temporal models.

This paper addresses three core questions:

- Can machine learning models effectively classify cell colony images at different timestamps?
- What insights are the networks deriving from the images to facilitate classification?
- How do sequential models compare to single-timestamp models in terms of performance?
- What temporal relationships can be discerned from the classification results provided by these models?

Through this exploration, our research highlights the significant potential of integrating computational methods with biological research, aiming to push the boundaries of current capabilities in modeling biological processes. The findings and methodologies developed from this study are expected to significantly advance computational techniques in biological research, offering new avenues for the study of morphogenesis and cell biology.

## III. Related Work

The landscape of cell image classification has undergone advancements in recent years, driven by the intersection of machine learning techniques and biomedical research. This cross-disciplinary area has sparked numerous studies, each exploring different methods to overcome cell image classification challenges.

Central to these endeavors has been the adoption of Convolutional Neural Networks (CNNs), which have become the cornerstone for cell image classification tasks [4]–[10]. CNNs have exhibited unparalleled efficacy in distilling and leveraging the intricate patterns present in cell imagery, fundamentally transforming our ability to interpret these visual data sources. Furthermore, the practice of Transfer Learning has further augmented the capabilities of CNNs in this domain [11]–[14]. Despite the successes of Convolutional Neural Networks (CNNs) and Transfer Learning in cell image classification, these approaches primarily excel in analyzing static images without considering the temporal dynamics inherent in cellular processes.

Our research explores the application of temporal models in image classification, spotlighting the innovative efforts that have advanced this field. The integration of Convolutional Neural Networks (CNNs) with Recurrent Neural Networks (RNN) [15]–[19] and the adoption of Transformer models for sequential image and video classification [20]– [23] exemplify the significant potential of sequential modeling in image classification tasks. A notable contribution by Karpathy et al. in the field of large-scale video classification using CNNs [15] particularly highlights the nuanced challenges encountered and the modest advancements achieved through incorporating spatio-temporal information into CNN architectures. This study reveals that, despite progress, the enhancements from embedding temporal dynamics within CNN frameworks tend to be more incremental than transformative.

Our study uniquely contributes by systematically comparing several temporal models in the context of cell image classification, contributing to the field of computational biology. We aim to identify models that best interpret temporal information in cell image sequences, thereby improving our understanding and predictive capabilities regarding cellular dynamics. This research not only challenges existing methodologies for temporal image classification but also opens new pathways for developmental biology research.

## IV. Methods

### A. Dataset

Our research utilizes a comprehensive dataset derived from the study conducted on human induced pluripotent stem cells (hiPSCs) to investigate the intrinsic cellular behaviors that guide morphogenesis during early lineage specification [3]. The dataset encompasses a collection of time-lapse images capturing the dynamic evolution of hiPSC colonies under various experimental conditions. The dataset serves as a valuable resource for developing and testing image classification algorithms, particularly those aimed at distinguishing between different states of cellular development. The temporal aspect of the dataset offers unique opportunities to apply and refine time-series analysis in the context of developmental biology.

The dataset comprises 78 unique samples, categorized based on the treatment conditions applied to the hiPSC colonies as follows:

1) WT: Wild Type/Untreated control samples

2) DS: Dual SMAD neuroectoderm protocol

3) BMP4: BMP4-induced trilineage protocol

4) CHIR: WNT activator CHIR

5) DS+CHIR: Addition of CHIR pre-treatment to the dual SMAD neuroectoderm

The distribution of samples over the 5 different categories is shown in table I. Figure 1 shows some sample images from the 5 classes. Each sample contains a sequence of 289 time-lapse images, corresponding to the 289 timestamps with a 5-minute interval across the 24-hour experimental duration, total to 22542 images, providing a rich temporal dimension to analyze cell behavior, morphological changes, and differentiation patterns.

**TABLE I:** Number of Samples for Each Category.

| Treatment | Number of Samples |
| --- | --- |
| WT | 12 |
| DS | 17 |
| BMP4 | 16 |
| CHIR | 15 |
| DS+CHIR | 17 |
| **Total** | **78** |

**Fig. 1:**
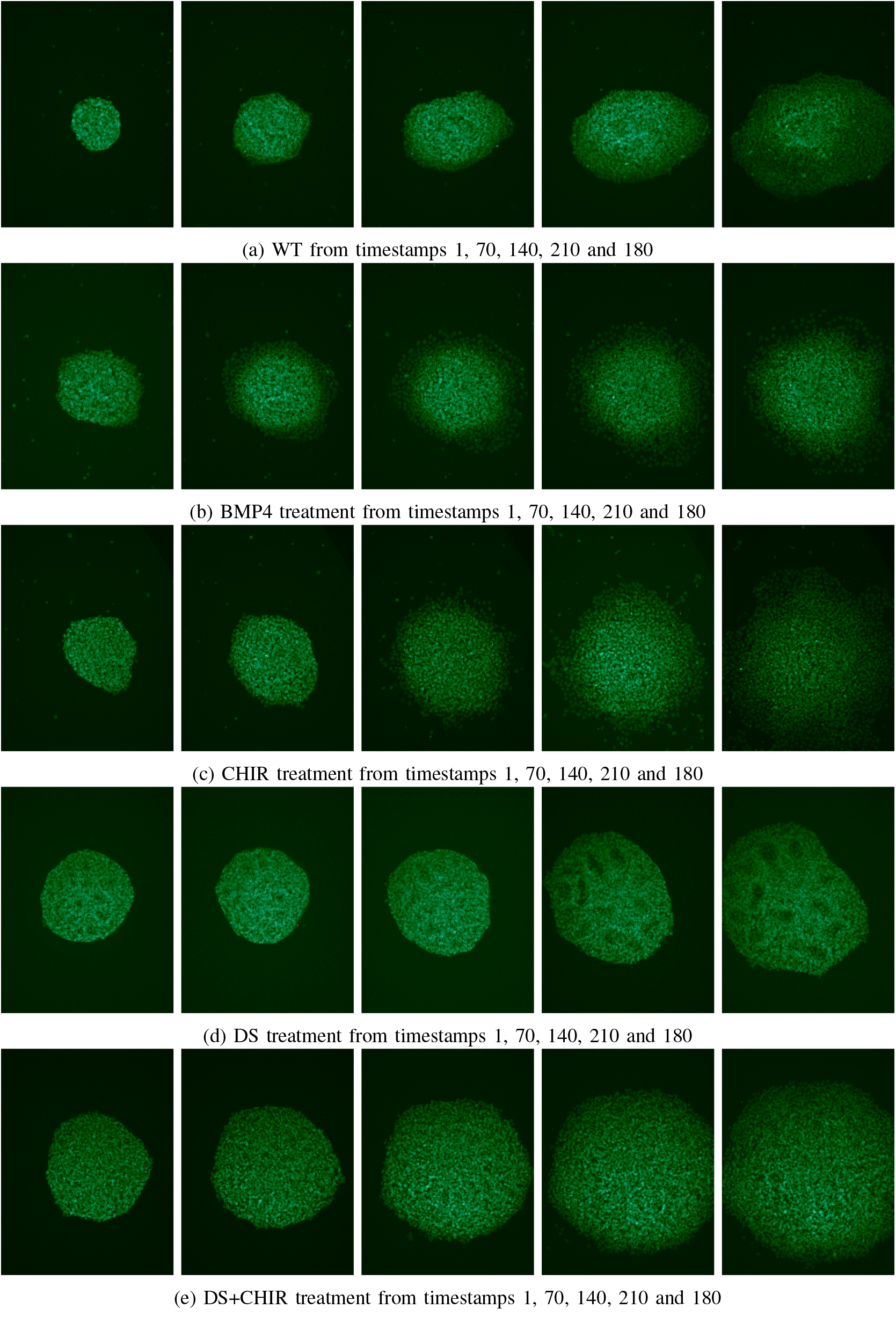
Sample images from dataset for the 5 treatments

### B. Non temporal Approach

In the non-temporal approach to cell image classification, each timestamped image of the same sample is considered an independent entity, disregarding temporal sequence. We utilized the power of Convolutional Neural Networks (CNNs) and transfer learning to develop separate classification models for each timestamp. Through this approach, we achieved promising results by training and testing with various pre-trained deep learning models. This approach not only establishes a benchmark for accuracy in identifying distinct cellular features but also lays the groundwork for conducting comparative analysis to aid in the exploration of cellular dynamics.

#### 1) Convolutional Neural Networks

Convolutional Neural Networks (CNNs) have become the baseline standard in image classification. CNN’s unique architecture is specifically good for capturing both the local details and global structure present within images [24]. A CNN typically consists of convolutional layers that apply various filters to the input images, pooling layers that reduce dimensionality, and fully connected layers that make the final classification decision [25].

Cell image classification with CNNs presents unique challenges, particularly due to the high variability in cell images and the often limited size of annotated datasets [26]. Cells can exhibit a wide range of appearances even within the same category, due to differences in staining, imaging conditions, and biological variability. Moreover, the scarcity of labeled data in cell imaging can hinder the ability of CNNs to learn effectively. We will utilize different methods in IV-B2, IV-B3 to overcome these obstacles and further optimize our results.

#### 2) Transfer Learning

At the core of transfer learning lies the assumption that features learned from one task or domain can be beneficial for solving related tasks or domains. Transfer learning in machine learning/deep learning references the process of applying the knowledge learned from a pre-trained machine learning model to a different but related problem [27]. The new problem can then be trained with less data and achieve better results with less computational power. The pretrained models are trained on large and comprehensive datasets, during which they adjust their internal parameters, such as weights and biases based on the data. In the context of image classification, pretrained models are trained on large image datasets such as ImageNet [28] under different architectures. Since the datasets are general in content and include rich categories of objects and features, the pretrained models thus can extract optimal features from all kinds of images.

Training deep learning models, particularly CNNs, often requires large datasets that are not always available in specialized scientific fields. Transfer learning addresses this by enabling models trained on small datasets to still perform effectively. Additionally, transfer learning significantly reduces the computational resources and training time required, making it a valuable approach for domains facing data scarcity and computational constraints. In our research, we employed transfer learning to train with several state-of-the-art CNN architectures, each known for its unique strengths in feature extraction and classification capabilities. These include:

- AlexNet [29]: As one of the pioneering architectures in deep learning, AlexNet laid the groundwork for CNNs in image classification.

Its architecture, though simpler compared to newer models, provides a solid starting point for transfer learning in our study.

- VGG [30]: Known for their deep architectures and excellent performance in capturing intricate textures and patterns, VGG models are highly effective in distinguishing subtle differences in images.
- GoogLeNet (Inception v1) [31] and Inception v3 [32]: These models introduce the concept of inception modules, allowing the network to adapt to various scales of image features.
- DenseNet [33]: Utilizes dense connections between layers to ensure maximum information flow, enhancing feature extraction capabilities for complex image structures.
- ResNet [34] and ResNeXt [35], including its variant Wide ResNeXt [36]: These architectures introduce residual connections to facilitate training of very deep networks, enabling them to learn high-level features without the vanishing gradient problem.

For each of these architectures, we initiated the transfer learning process by replacing the final classification layer with a new layer tailored to our specific task — classifying 5 types of cell images. During the process, the weights of the layers before the final classification layer are frozen (fig. 2). This approach of transfer learning can be referred to as feature extractor (as opposed to fine-tuning from scratch) which takes the advantage of minimizing computational costs [37] for faster training.

**Fig. 2:**
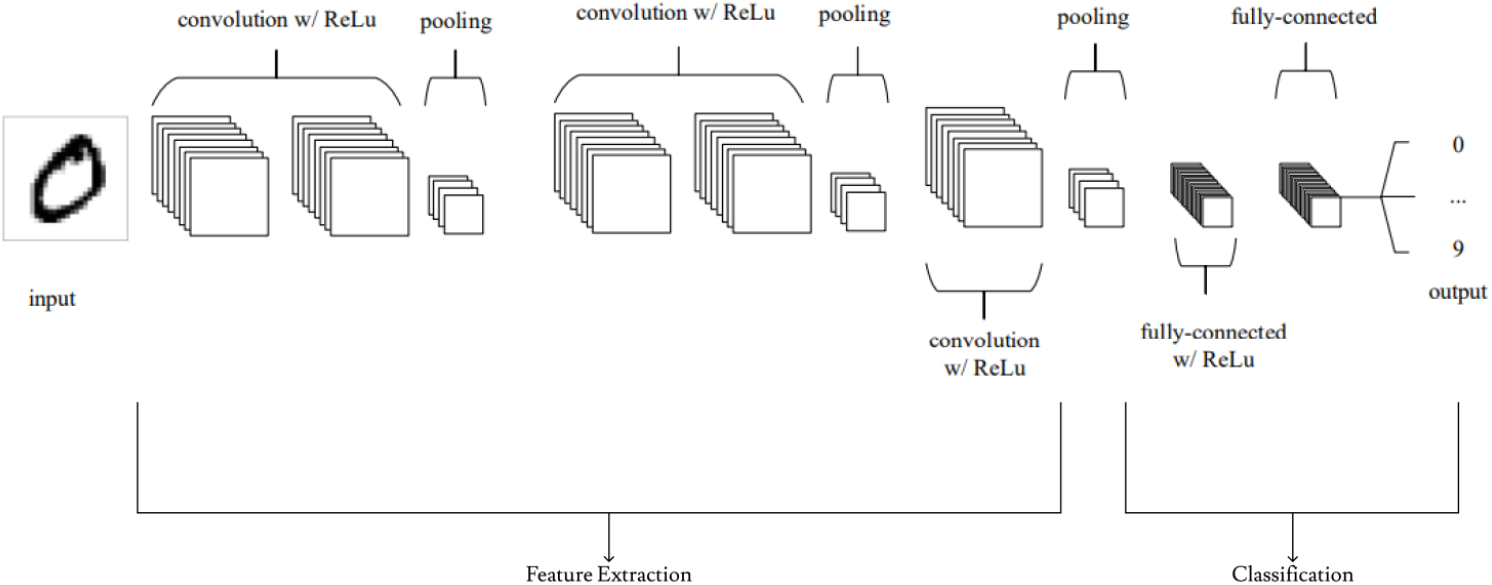
Transfer Learning with a common CNN architecture

Through transfer learning, we aim to capitalize on the knowledge embedded in pre-trained models and adapt it to the specific challenges of cell image classification, ultimately improving the accuracy and efficiency of cell image classification.

#### 3) Data Enrichment

Data enrichment, a crucial step in preparing datasets for deep learning models, involves augmenting the original dataset to enhance model training and performance. In the context of our cell image classification project, we implemented data enrichment by rotating images in our dataset. This technique not only increases the volume of data available for training but also introduces a variety of perspectives, enabling the model to learn more generalized features of the cell images.

For our original dataset, initially comprising 78 images split into training, validation, and testing sets with a ratio of 8:1:1, resulting in only 62 samples for training, 8 for validation and 8 for testing. We applied a simple yet effective data augmentation strategy: rotating each image in four directions. By rotating each image in four orientations (0°, 90°, 180°, and 270°), we effectively quadrupled our dataset size. The new distribution consisted of 249 training samples, 31 validation samples, and 32 testing samples with an 8:1:1 ratio and 218 training samples, 47 validation samples, and 47 testing samples with a 7:1.5:1.5 ratio. This enrichment process ensured that our model was not just learning from a limited set of images but was exposed to a broader variation of cell orientations, crucial for improving the model’s ability to accurately classify cell images under different conditions.

By augmenting the number of images, we mitigate the risk of overfitting, a common challenge when training deep learning models with limited data [38]. Introducing varied orientations of the same cell images helps the model learn more comprehensive features, enhancing its ability to generalize from the training data to new, unseen images. Most importantly, with the enriched dataset, we were able to achieve better model accuracy across all base models used in transfer learning.

### C. Temporal Approaches

In this section, we introduce three temporal approaches tailored for our cell image classification task: Long Short Term Memory Networks (LSTM), R-Transformer with feature extraction and Video Vision Transformer(ViVit). Our hypothesis posits that these temporal methodologies can proficiently capture the intricate dynamics present in our sequential images and thus can outperform the models trained for separate timestamps.

#### 1) Feature Extraction

Extracting the features from images is an important step in many image processing tasks as it extracts meaningful information and characteristics from images [39]. In the general CNN architecture as shown in fig. 2, all the layers before the classification step (fully connected layer) are part of feature extraction, aiming to identify distinctive patterns, structures, or attributes within an image. The importance of feature extraction lies in its ability to transform high-dimensional image data into a more compact and informative representation, facilitating more efficient and effective analysis [39].

In our experiments, we aim to leverage the features extracted from pre-trained models which are more compact and informative representations of the input images. We then group sequences of extracted features from images of the same sample together and input them into our LSTM and Rtransformer model.

### 2) RNN/LSTM + Feature Extraction

Recurrent Neural Networks (RNN) and their advanced variant, Long Short Term Memory Networks (LSTM), are designed to handle sequential data with the capability to capture dependencies at different time scales. Their architectures are shown in fig. 3. RNNs are adept at modeling short-term dependencies due to their recurrent structure, which processes sequences one element at a time, maintaining a ‘memory’ of previous inputs [41]. LSTMs extend this capability by incorporating mechanisms—such as input, output, and forget gates—that allow the network to retain or discard information over longer periods [42]. This makes LSTMs particularly suited for tasks where understanding both immediate and extended context within the data is crucial.

**Fig. 3:**
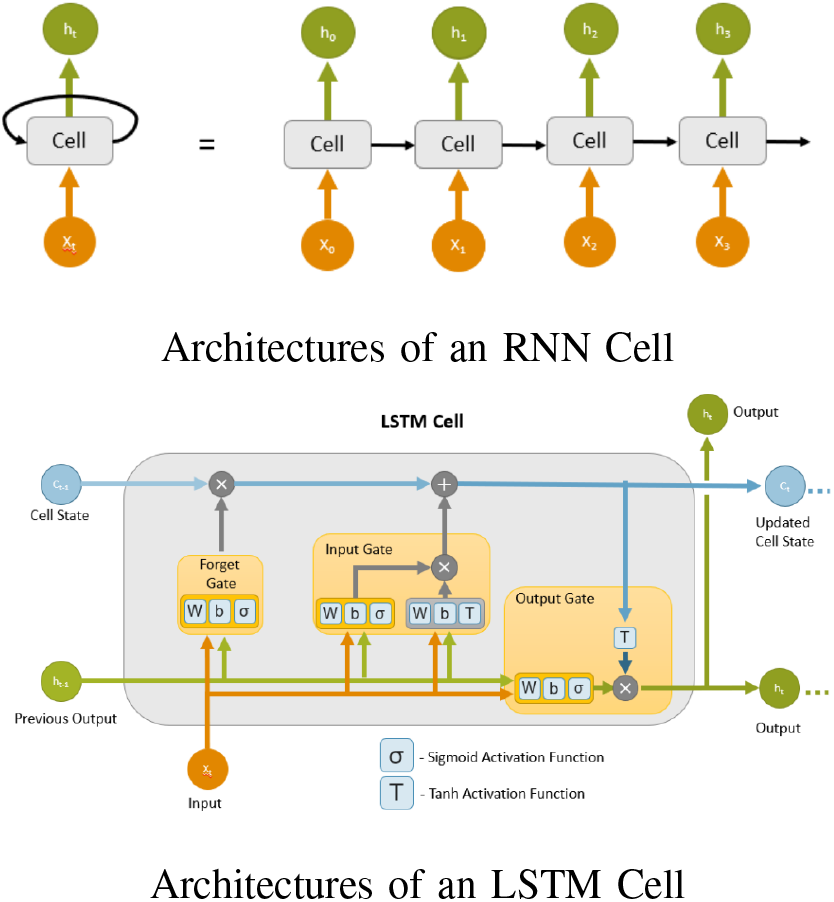
Architectures of RNN and LSTM Cell [40]

In our approach, we utilize LSTMs in conjunction with feature extraction from pre-trained CNN models. By doing so, we aim to harness the detailed, high-level features extracted from each image frame, presenting these as a sequence to the LSTM. This allows the LSTM to analyze the temporal evolution of these features across the sequence of images.

### 3) R-Transformer + Feature Extraction

While RNNs and their variants have laid the groundwork for sequence modeling, their limitations in processing long-term dependencies and parallelization have led to the development of transformer models. Transformers revolutionize sequence modeling through their attention mechanisms, enabling the model to weigh the importance of different parts of the input data without the constraints of sequential processing [43]. However, transformers typically require significant computational resources and may struggle with capturing localized sequence patterns due to their global attention mechanism.

The R-transformer architecture combines the strengths of RNNs and transformers, addressing both local and global modeling of sequence data [44]. Fig. 4 shows one R-transformer layer containing localRNNs that process local sequence pattern, multi-head attention networks for global context understandings and Position-wise feedforward networks that transform the features non-linearly. This hybrid model is particularly effective in our context, where capturing the detailed evolution at each timestamp and the overarching trends across the sequence is crucial for accurate classification.

**Fig. 4:**
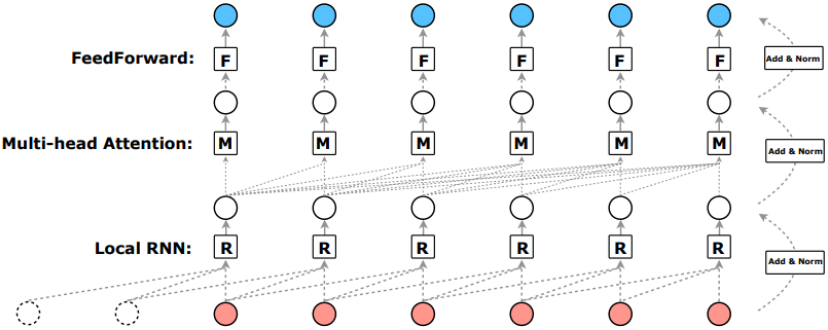
Architectures of one layer of R-Transformer [43]

### 4) ViVit Transformer

The ViVit Transformer architecture represents a leap forward in processing spatio-temporal data, such as video sequences, by applying the principles of transformers to capture both spatial and temporal dynamics [20]. Unlike traditional CNNs that may require complex architectures to handle time series data, ViVit simplifies this process by treating the sequence of images as spatio-temporal tokens as shown in fig. 5. This approach enables the model to directly learn the intricate patterns of change within the data, making it an excellent candidate for our task of classifying cell images across different timestamps.

**Fig. 5:**
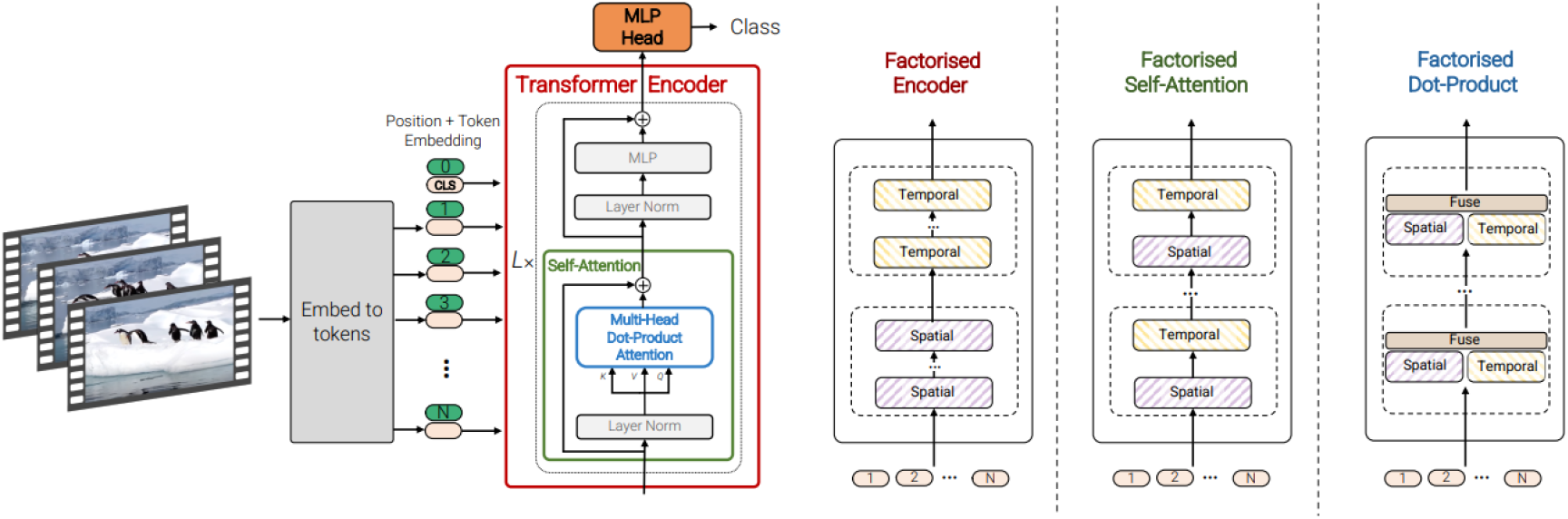
Architectures of ViVit [20]

Our implementation of these temporal models—LSTM, R-Transformer, and ViVit—aims to leverage their unique strengths in capturing the temporal dynamics present in sequential cell images. By comparing their performance with the single timestamp models, we seek to identify the most effective methods for understanding the complex evolution of cell colonies, potentially setting new standards for temporal analysis in cell image classification.

## V. Results and Comparisons

### A. Non-Temporal Approach Results

#### 1) Comparison of Transfer Learning Models Overall Performance

Table II and III presents a detailed comparison of the average accuracy and training time for a range of pre-trained models employed in the classification of cell colony images across 289 timestamps. These architectures were consistently trained using identical epochs and hyperparameters to ensure a fair comparison.

**TABLE II:** Average Accuracy and Training Time of Transfer Learning Models with training/validation/test ratio 8:1:1.

| Model | Without Data Enrichment | With Data Enrichment | Training Time/Timestamp (s) |
| --- | --- | --- | --- |
| AlexNet | 87.0675 | **95.5125** | **2** |
| VGG16 | 89.2734 | **96.3884** | **18** |
| GoogleNet | 88.7111 | 95.4477 | 8 |
| Inception_v3 | 87.7163 | 94.7664 | 14 |
| ResNet18 | 89.5761 | 93.4256 | 5 |
| ResNet50 | 86.4187 | 94.5609 | 11 |
| DenseNet | 91.2630 | **96.3020** | **23** |
| ResNeXt | 89.0571 | 94.3555 | 11 |
| Wide ResNeXt | 84.3858 | 93.9446 | 16 |

**TABLE III:** Average Accuracy and Training Time of Transfer Learning Models with training/validation/test ratio 7:1.5:1.5.

| Model | Without Data Enrichment | With Data Enrichment |
| --- | --- | --- |
| AlexNet | 88.2353 | 89.1188 |
| VGG16 | 87.5144 | 90.4734 |
| GoogleNet | 89.3022 | 94.0587 |
| Inception_v3 | 89.3022 | **94.3164** |
| ResNet18 | 84.2849 | 93.0796 |
| ResNet50 | 87.6298 | 91.6955 |
| DenseNet | 91.7820 | **95.7079** |
| ResNeXt | 88.6678 | 91.7249 |
| Wide ResNeXt | 85.1211 | 93.1090 |

It is evident that all models experienced significant performance improvement when data enrichment techniques were applied, underscoring the importance of augmenting training data to enhance model accuracy, particularly in scenarios with small datasets.

DenseNet emerged as the leading model in terms of accuracy after data enrichment, achieving remarkable performance with an average accuracy of 96.3020% and 95.7079% for test ratios of 0.2 and 0.3, respectively. Inception v3 also performed admirably with 94.7664% and 94.3164%. Notably, these two models, while providing excellent accuracy, also required the most substantial computational resources, as reflected in their longer training times of 23 and 14 seconds per timestamp, respectively. GoogleNet, on the other hand, demonstrated a consistent performance with accuracies of 95.4477% and 94.0587% across both test ratios, combined with a relatively quick training time of 8 seconds per timestamp.

Interestingly, while VGG16’s accuracy was notably high with a training ratio of 0.8 at 96.3884%, its performance declined to 90.4734% when the dataset was divided with a training ratio of 0.7. This drop highlights VGG16’s sensitivity to the size of the training set, emphasizing the need for ample data to maintain its high classification accuracy.

In summary, while DenseNet and Inception v3 offer the highest accuracies, their computational demands are significant. GoogleNet presents a compelling alternative, offering a balanced compromise between accuracy and speed. VGG16’s variable performance across different dataset sizes further highlights the importance of selecting models based on specific project requirements and data availability.

### 2) Comparison of Model Performance Across Timestamps

The examination of model performance across various timestamps reveals critical insights into the dynamic behavior and evolution of cell colonies in response to treatments over time. By analyzing the accuracy plots for the models across all timestamps (fig. 6), we can infer the following patterns:

**Fig. 6:**
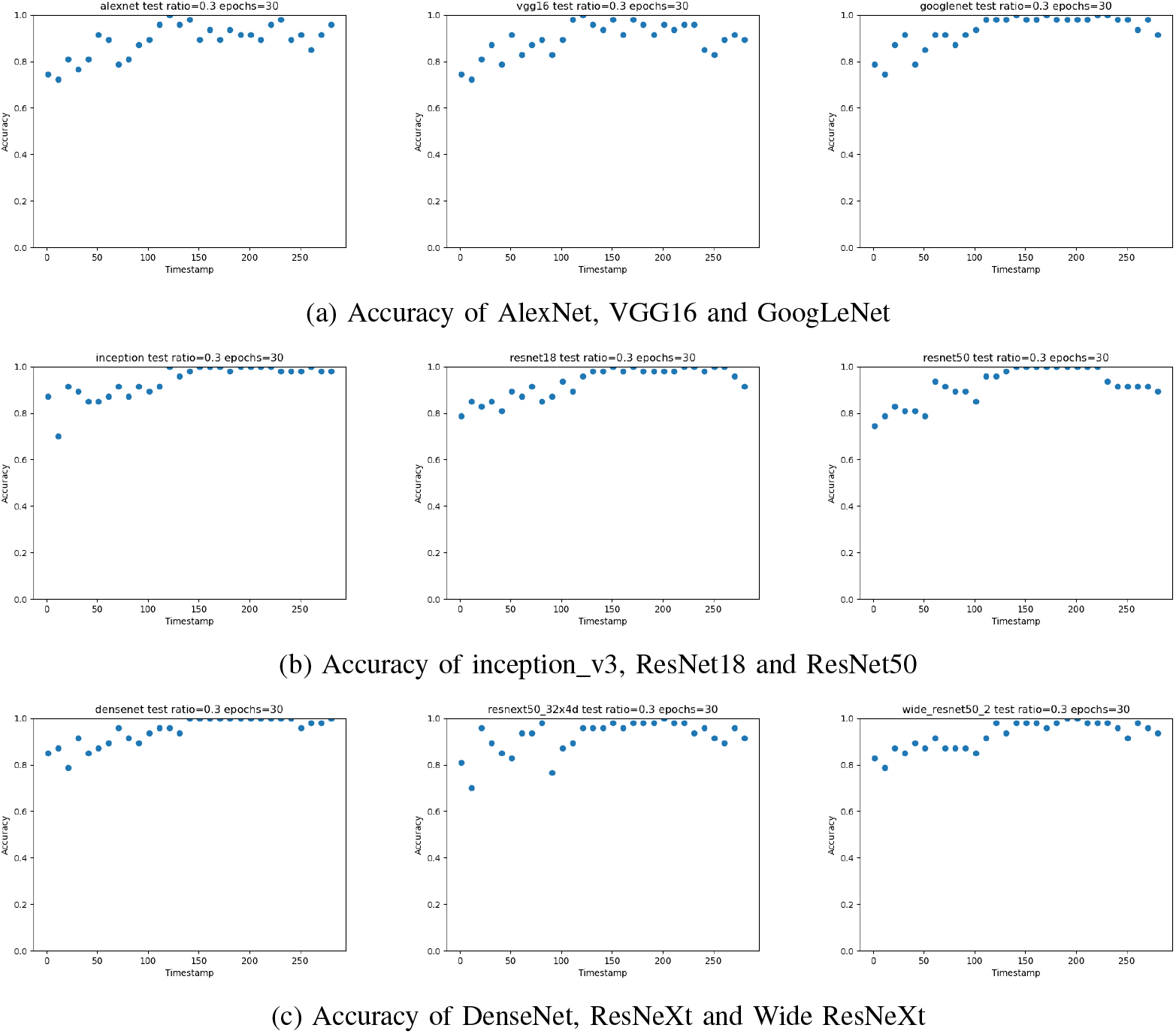
Accuracy of Transfer Learning Models

**Fig. 7:**
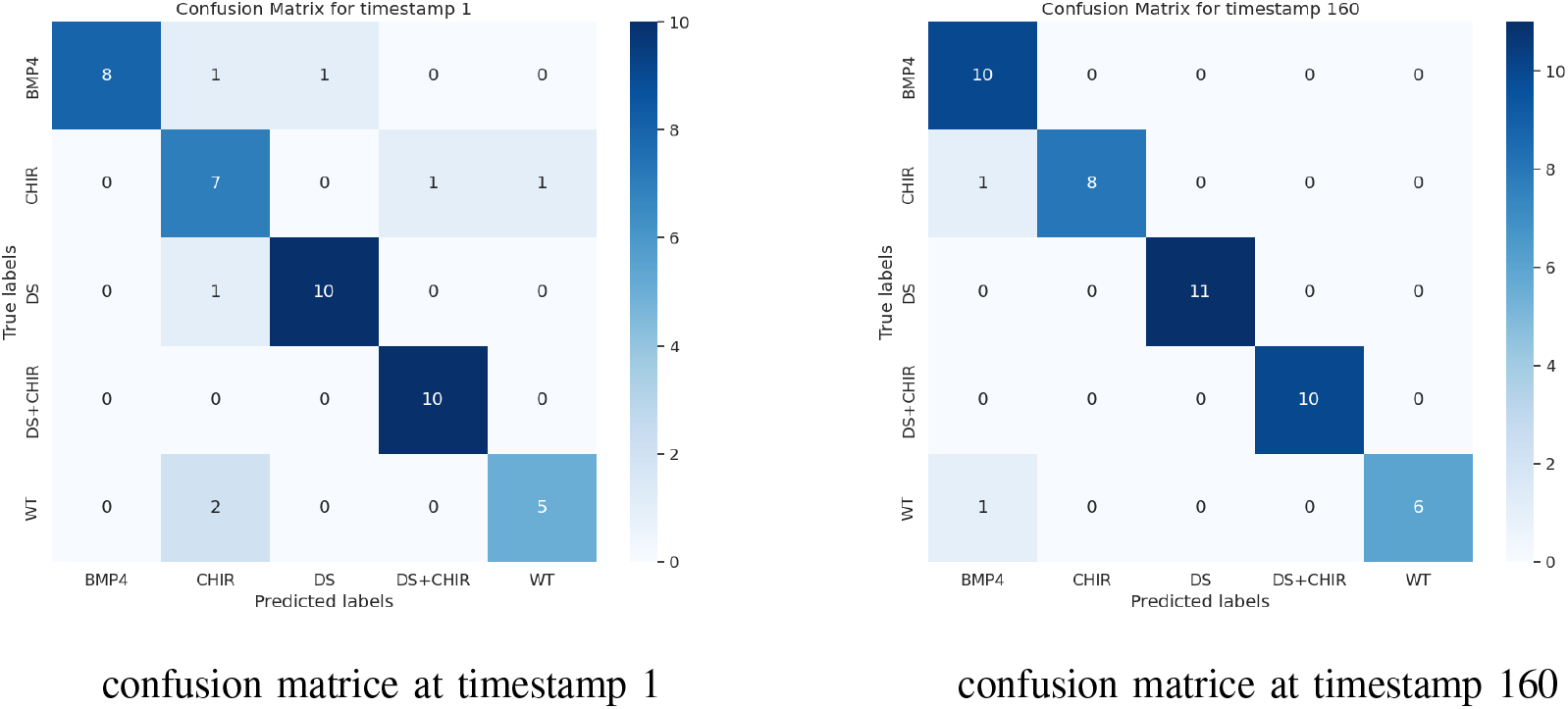
Example of DenseNet model’s confusion matrices

#### a) Early Stage Performance

Initially, at the first timestamp, all models show lower accuracies, suggesting that the cell colonies have yet to exhibit significant divergences. Interestingly, although lower, the accuracy is not minimal, implying that the models must be picking up on early, nuanced differences post-treatment. This observation opens promising pathways for future research to explore the early cellular responses to treatments and the subtle morphological changes or features these models are detecting, potentially offering insights that may not be easily observable by human analysts. .

#### b) Trend Over Time

As expected, there is a noticeable improvement in model accuracy over time, reflecting the increasing distinctiveness of cell features as colonies evolve. This trend is particularly evident in models like inception v3, DenseNet and googlenet, which demonstrate consistently high performance. The increasing accuracy over time aligns with the hypothesis that as cellular features become more defined, they are easier for models to classify correctly.

#### c) Ending Stage Performance

Across all models, a decrease in accuracy at later timestamps becomes evident, particularly in models like VGG16, ResNet18, ResNet50, and ResNeXt. This trend suggests a convergence in the models’ ability to differentiate between cell states as the experiment progresses. Notably, this fall may indicate that the cell colonies have reached a developmental stage characterized by minimal changes in their features or that these features have become increasingly challenging for the models to distinguish. Moreover, this phenomenon could imply that the cells have developed into similar forms by these later stages, making discrimination more difficult for automated analysis. This observation prompts further investigation into biological aspects—such as pinpointing the stages at which cellular features stabilize or become visually indistinguishable.

### 3) Analysis of Misclassifications

The confusion matrices for the models at various timestamps provide a deep dive into the classification performance and the evolution of the model’s predictive capabilities over time, as illustrated in examples shown in 7. Additionally, Table IV, which compiles statistics of misclassified pairs across all models and timestamps, helps identify specific patterns:

**TABLE IV:** Misclassified Classes from timestamps 1, 40, 80, 120, 160, 200, 240, 280, 289 from all models. (Note: that the total column represents the total number for all timestamps instead of the shown ones, thus it is not the sum of the row)

| Misclassified Pair/Count | Timestamps |  |  |  |  |  |  |  |  | Total |
| --- | --- | --- | --- | --- | --- | --- | --- | --- | --- | --- |
|  | 1 | 40 | 80 | 120 | 160 | 200 | 240 | 280 | 289 |  |
| WT vs. BMP4 | 11 | 7 | 13 | 0 | 2 | 0 | 0 | 0 | 0 | 1133 |
| WT vs. CHIR | 14 | 14 | 8 | 1 | 0 | 0 | 0 | 1 | 2 | 1301 |
| WT vs. DS | 0 | 0 | 0 | 2 | 0 | 0 | 2 | 0 | 1 | 136 |
| WT vs. DS+CHIR | 0 | 0 | 0 | 0 | 1 | 1 | 7 | 10 | 9 | 459 |
| BMP4 vs. CHIR | 14 | 15 | 7 | 4 | 8 | 13 | 8 | 5 | 8 | 2863 |
| BMP4 vs. DS | 1 | 2 | 0 | 0 | 0 | 0 | 0 | 0 | 0 | 40 |
| BMP4 vs. DS+CHIR | 3 | 0 | 0 | 0 | 0 | 0 | 4 | 5 | 0 | 544 |
| CHIR vs. DS | 6 | 1 | 0 | 0 | 0 | 0 | 0 | 0 | 0 | 127 |
| CHIR vs. DS+CHIR | 14 | 15 | 17 | 4 | 0 | 0 | 1 | 8 | 7 | 2248 |
| DS vs. DS+CHIR | 14 | 2 | 0 | 0 | 1 | 0 | 0 | 1 | 2 | 211 |
| **Total** | 77 | 56 | 45 | 11 | 12 | 14 | 22 | 30 | 29 | - |

Misclassifications WT vs. BMP4 and WT vs. CHIR are notably high in the initial timestamps but significantly decrease or virtually disappear in later timestamps. This trend suggests that BMP4 and CHIR treatments induce slower initial changes in cell morphology, making these cells more challenging to distinguish from untreated WT cells early on. However, as the experiment progresses, the distinctive features of BMP4 and CHIR-treated cells become more pronounced, leading to improved classification accuracy.

Conversely, DS treatment is associated with the fewest misclassifications among other classes, likely due to the distinct morphological characteristics it induces in cells, such as shallow spots, which are readily identifiable in sample images (see Figure 1).

Interestingly, while pairs like BMP4 vs. CHIR and CHIR vs. DS+CHIR continue to be confused across nearly all timestamps, indicating their similarity, BMP4 vs. DS+CHIR pairs show a lower rate of misclassification. This suggests that the likelihood of misclassifying certain treatment pairs does not necessarily follow a pattern of transitivity, highlighting the complexity of cellular responses to different treatments.

Moreover, it’s crucial to note that the highest incidence of misclassifications occurs at the first timestamp across most pairs. Yet, specific pairs such as WT vs. DS and WT vs. DS+CHIR show zero misclassifications even from the outset. This zero misclassification rate for such pairs from the first timestamp underscores the immediate and distinct impact of DS and DS+CHIR treatments on cell morphology, making these cells instantly recognizable compared to WT, even at the earliest stage of the experiment. This clear delineation from the beginning suggests a rapid onset of morphological changes that are readily captured by the models, contrasting with the more gradual evolution observed with BMP4 and CHIR treatments.

### 4) Heatmaps Analysis

The analysis of modelgenerated heatmaps offers a window into the neural network’s decision-making process and allows us to understand the features deemed significant for classifying cell images. This comparison between the features the model focuses on and human precepts is crucial for validating the model’s effectiveness in uncovering subtle biological features and whether it includes more details than the human eye.

In fig. 8, we show some examples of heatmaps generated by DenseNet and VGG models. While DenseNet leads in accuracy, the VGG-generated heatmaps align more closely with human interpretations of cellular characteristics. For example, VGG’s attention to the distinctive holes in DS-treated cell colonies and the fading edges indicative of CHIR treatment mirrors the human eye’s cues for these conditions. Despite its marginally lower accuracy, this alignment suggests VGG’s potential to provide interpretable insights into treatment effects on cells presented in the images.

**Fig. 8:**
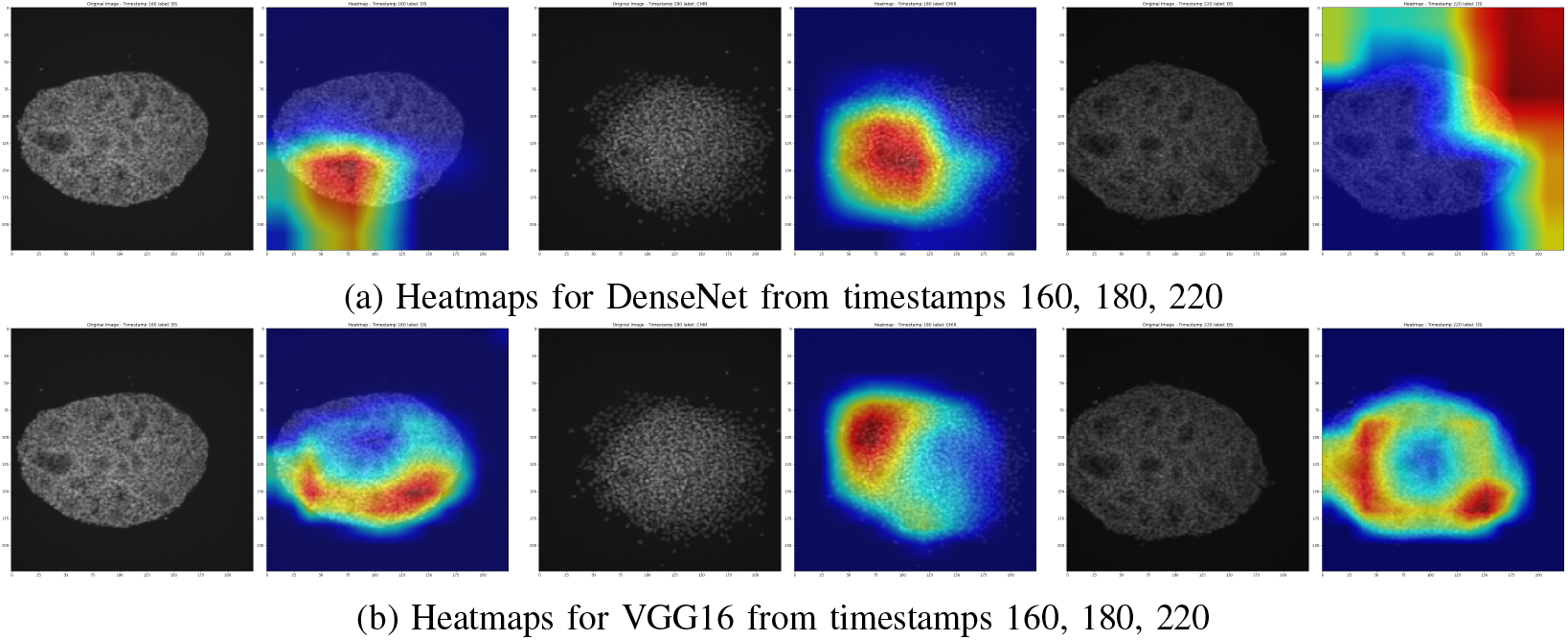
DenseNet vs. VGG heatmaps on the same set of correctly predicted images

Moreover, the discrepancy between DenseNet’s classification success and its heatmap focus areas presents a curious case for deeper analysis. Despite DenseNet’s high accuracy, its heatmaps do not always emphasize the same key features that VGG does, which are readily identifiable by human observers as indicative of specific treatments. This divergence raises intriguing questions about the underlying mechanisms of DenseNet’s learning process and its ability to classify images accurately without apparently prioritizing the same visually distinctive features. It suggests that DenseNet might be leveraging more subtle, perhaps less intuitive aspects of the images that are not immediately obvious to human analysts.

The frequent misclassifications between BMP4 and CHIR treatments, as evidenced in our confusion matrices, further underscore the challenges in distinguishing these conditions. Heatmaps from the VGG model, particularly Fig. 9, illustrate a dynamic focus: early attention to the cell’s core morphological features gradually expands to include peripheral or adjacent areas. This shift may reflect the model’s adaptation to more nuanced, time-evolving cellular responses not immediately apparent in the initial stages. These observations suggest a layered complexity in how different treatments manifest in cellular morphology over time. The initial model focuses on central cell features, followed by a broader consideration of surrounding areas, hints at a temporal dimension in cellular response patterns that models begin to recognize and attempt to decode.

**Fig. 9:**
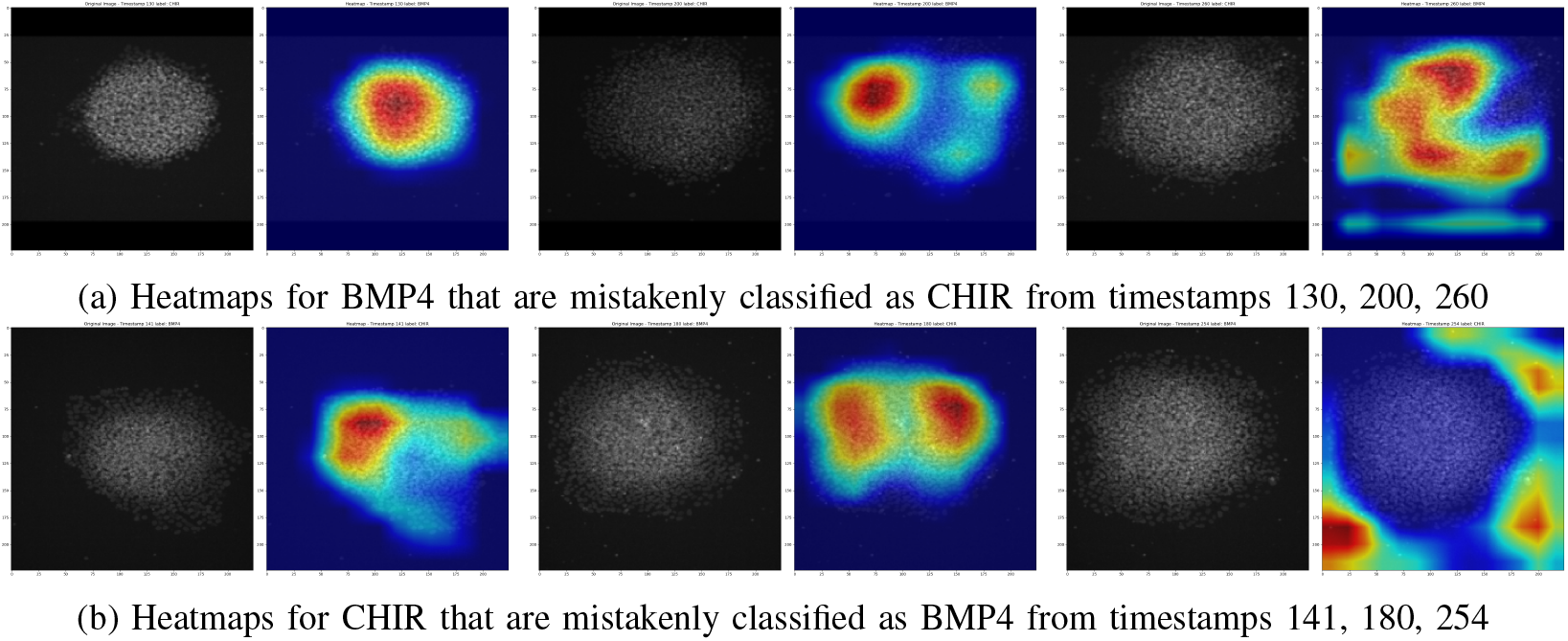
Heatmaps for misclassification pair BMP4 vs. CHIR

#### B. Temporal Approach Results

Our experimental setup was designed to rigorously evaluate the performance of both nontemporal and temporal models in classifying cell colony images across a series of developmental stages represented by timestamps. To this end, we utilized a set of pre-trained models for non-temporal classification tasks and adapted temporal models—LSTM, R-Transformer, and ViVit—for a comparative analysis. For LSTM and R-Transformer, we extracted features and input a sequence of features for the sequence of images to the network, whereas ViVit directly processed sequences of images as input. For LSTM and R-Transformer, we opted for feature extraction using VGG16, a decision underpinned by experimental observations that VGG16 generally outperforms DenseNet as a feature extractor. This preference is further supported by heatmap analyses V-A4, which suggest VGG16’s superior capability in highlighting relevant features for classification tasks.

Tables V and VI present a comparison of model accuracies across various timestamp ranges of cell colonies. For non-temporal models (DenseNet and VGG16), the accuracies reported are the average performances for each of the models within each specified timestamp range. In contrast, for temporal models, including LSTM, R-Transformer, and ViVit, we have a single model for each timestamp range, to ensure a balanced comparison, the accuracies for these models are averaged over multiple runs. For enhanced clarity, the highest accuracy within each timestamp range is highlighted in the tables.

**TABLE V:** Comparison of Average Accuracy Over Timestamp Ranges for VGG16, LSTM and RTransformer and ViVit.

| Timestamp Range/<br>Model | Non Temporal<br>VGG16 | Temporal |  |  |
| --- | --- | --- | --- | --- |
|  |  | LSTM | R-Transformer | ViVit |
| 1-20 | **86.9048** | 85.1063 | 83.6879 | 82.2695 |
| 21-40 | 83.3333 | 82.2695 | **84.3972** | 82.9787 |
| 41-60 | 80.9524 | 83.6879 | 85.8156 | **88.6525** |
| 61-80 | 83.3333 | 88.6525 | **87.9433** | 87.2340 |
| 81-100 | 82.1429 | 87.2340 | 88.6525 | **89.3617** |
| 101-120 | 84.1270 | **95.0355** | 92.9078 | 87.2340 |
| 121-140 | 90.8730 | **97.8723** | 94.3262 | 93.6170 |
| 141-160 | 83.3333 | **91.4894** | 90.780 | 90.0709 |
| 161-180 | 84.5238 | 90.7801 | **95.7447** | 89.3617 |
| 181-200 | 89.2857 | **90.0709** | **90.0709** | 87.2340 |
| 201-221 | **95.2381** | 95.0354 | 93.6170 | 87.9433 |
| 221-240 | **94.0476** | 88.6525 | 92.1986 | 90.7801 |
| 241-261 | 96.0317 | 87.2340 | 87.2340 | **97.8723** |
| 261-281 | **92.4603** | 85.8156 | 90.7801 | 88.652 |
| 281-289 | **85.1852** | 82.9787 | 85.1064 | 85.1064 |
| 130-220 | 88.7363 | 93.6170 | **95.0354** | 94.3262 |

**TABLE 6:** Comparison of Average Accuracy Over Timestamp Ranges for DenseNet, LSTM, RTransformer and ViVit.

| Timestamp Range/<br>Model | Non Temporal<br>DenseNet | Temporal |  |  |
| --- | --- | --- | --- | --- |
|  |  | LSTM | R-Transformer | ViVit |
| 1-20 | 82.5397 | **85.1063** | 83.6879 | 82.2695 |
| 21-40 | 82.9365 | 82.2695 | **84.3972** | 82.9787 |
| 41-60 | 79.3651 | 83.6879 | 85.8156 | **88.6525** |
| 61-80 | **90.8730** | 88.6525 | 87.9433 | 87.2340 |
| 81-100 | 88.4921 | 87.2340 | 88.6525 | **89.3617** |
| 101-120 | 90.4762 | **95.0355** | 92.9078 | 87.2340 |
| 121-140 | 94.0476 | **97.8723** | 94.3262 | 93.6170 |
| 141-160 | **93.2540** | 91.4894 | 90.780 | 90.0709 |
| 161-180 | 95.2381 | 90.7801 | **95.7447** | 89.3617 |
| 181-200 | **97.2222** | 90.0709 | 90.0709 | 87.2340 |
| 201-221 | **95.6349** | 95.0354 | 93.6170 | 87.9433 |
| 221-240 | **96.8254** | 88.6525 | 92.1986 | 90.7801 |
| 241-261 | **98.8095** | 87.2340 | 87.2340 | 97.8723 |
| 261-281 | **96.8254** | 85.8156 | 90.7801 | 88.6525 |
| 281-289 | **94.4444** | 82.9787 | 85.1064 | 85.1064 |
| 130-220 | **95.2380** | 93.6170 | 95.0354 | 94.3262 |

### 1) Temporal Models Exhibit Early Strength

Analysis of average accuracies over timestamp ranges reveals a distinctive pattern: temporal models generally achieve higher accuracy in the earlier stages of cell development (Tables V and VI). For example, in the initial 1-20 timestamp range, LSTM and R-Transformer models demonstrate a notable advantage over their non-temporal counterparts. This suggests that temporal models are particularly adept at capturing the subtle dynamics and changes occurring in the early stages of cell development.

### 2) Non-Temporal Models Gain in Later Stages

As cell colonies progress through their developmental stages, the advantage of temporal models diminishes. In later timestamp ranges, non-temporal models, particularly DenseNet, consistently outperform temporal approaches. This shift could be attributed to the evolving complexity of cellular features, which become more pronounced and easier for non-temporal models to discern individually. DenseNet, the best-performing single model, showcases superior overall accuracy across the majority of timestamp ranges, affirming its robustness in identifying diverse cellular states as they become more distinct.

Furthermore, an intriguing pattern emerged during our analysis, particularly with temporal models such as R-Transformer and ViVit. These models consistently identified BMP4 vs. CHIR treatments as their sole significant challenge, leading to misclassifications. As illustrated in the confusion matrix in Fig. 10, this specific pairing was the only one that consistently confused the models across most timestamp ranges. This unique misclassification points to a pronounced similarity between BMP4 and CHIR treatments, which, notably, was not as perplexing for non-temporal models like DenseNet. DenseNet’s ability to accurately distinguish between these two treatments in later timestamps—where temporal models continued to struggle—highlights a discernible gap in the models’ capabilities. This singular misclassification challenge raises pivotal questions about the distinctive features that DenseNet may be leveraging to achieve its classification success, underscoring a critical area for future research to refine the sensitivity of temporal models to the nuanced differences between such closely related treatments.

**Fig. 10:**
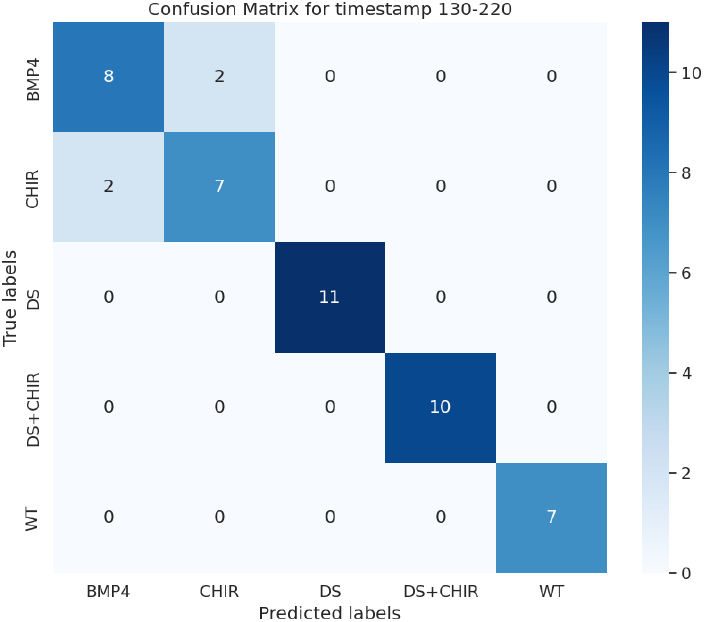
An example of a confusion matrix for timestamp ranges 130-220 for ViVit

## VI. Discussion and Future Directions

This study’s exploration into the classification of cell colony images across various developmental stages has underscored the nuanced capabilities of both non-temporal and temporal models. The differential performance of these models, particularly in early versus later stages of cell development, highlights the complexity of cellular evolution and the dynamic nature of biological processes.

Notably, our findings reveal a distinct advantage of temporal models in capturing the early dynamics of cell development, an area ripe for further exploration. This early-stage sensitivity could be pivotal in applications requiring early detection of cellular changes, such as in disease diagnosis or monitoring the efficacy of treatments.

In our analysis, DenseNet’s high classification accuracy presents a paradox when juxtaposed with its heatmap analysis, which does not consistently align with human visual interpretations. This discrepancy raises critical questions about the nature of features DenseNet leverages for its predictions. Unlike human observers who may rely on recognizable morphological traits, DenseNet might be identifying and utilizing more abstract or subtle features not immediately apparent to researchers. The consistent misclassification of BMP4 and CHIR treatments by temporal models, yet their clear distinction by DenseNet in later stages, raises important questions about the models’ feature prioritization and learning mechanisms. Future studies should aim to delve into the interpretability of these models, potentially through techniques like layer-wise relevance propagation or feature visualization, to uncover the underlying features that drive model decisions.

## VII. Conclusions

Our comprehensive analysis has demonstrated the potential of machine learning techniques, both nontemporal and temporal, to significantly advance our understanding of cellular dynamics. By leveraging state-of-the-art models and innovative sequence modeling techniques, we have provided new insights into the behavior of cell colonies over time, contributing to the broader field of computational biology.

Future research should focus on enhancing the interpretability of these models, exploring hybrid approaches that combine the strengths of both nontemporal and temporal models, and extending the application of these findings to practical biological and medical challenges. The journey towards fully understanding cellular evolution is complex and multifaceted, and this study represents a significant step forward in that ongoing exploration.

